# Intracellular spirochete integrates into a chemosynthetic micro-consortium of a deep-sea vent mussel

**DOI:** 10.64898/2026.11.17.763021

**Authors:** Tetsuro Ikuta, Yoshihiro Takaki, Ayu Yuasa, Yukari Yoshida-Takashima, Miwako Tsuda

## Abstract

Animal-microbe interactions comprise a diverse functional landscape, spanning pathogenic to mutualistic outcomes across extracellular and intracellular niches. Within this context, spirochetes are predominantly documented as extracellular associates or transient pathogens, leaving stable, functional endosymbiosis within animal cells unverified. Here, we report the discovery of an intracellular consortium comprising a spirochete and a primary sulfur-oxidizing bacterium within gill bacteriocytes of the deep-sea hydrothermal vent mussel *Gigantidas horikoshii*. Combining three-dimensional ultrastructural imaging, spatial histology, and multi-omics analyses, we characterize the intracellular spatial architecture and metabolic integration of this system. Both bacteria co-reside within a closed, membrane-bound network of interconnected chambers, where the spirochete is distributed across compartments and occasionally extends across host cell boundaries. Multi-omics profiling uncovers metabolic cooperation between the two bacteria: while the sulfur-oxidizing bacterium provides autotrophic primary production, the spirochete actively transcribes pathways for reactive oxygen species scavenging and organic acid assimilation, likely mitigating localized oxidative stress and acidosis. Coupled with its motility machinery and an expanded repertoire of host-interaction factors, these physiological traits suggest that the spirochete acts as an active stabilizer of the shared intracellular microenvironment. Together, these findings reveal that spirochetes can establish a stable, functional micro-consortium with chemosynthetic bacteria within animal cells, refining the recognized landscape of animal-microbe interactions and illuminating mechanisms linking mutualistic endosymbiosis and intracellular infections.

## Introduction

Interactions between animal hosts and microbes comprise a diverse and dynamic functional landscape (1, 2), in which pathogenic outcomes typically manifest as host tissue damage, whereas mutualistic partnerships promote host success through benefits such as nutritional support, ecological niche expansion, and evolutionary diversification (1, 3). Among these interactions, nutritional symbiosis represents a primary paradigm, where microbial partners synthesize essential nutrients to fuel host metabolism (4). The most pronounced form of this strategy occurs in partnerships with autotrophic microorganisms, which sustain host nutrition largely through primary production (5, 6). While binary host-microbe associations have served as classical models to dissect these phenomena (2, 7), natural symbioses increasingly reveal multipartite assemblages functioning as consortia, where distinct microbial associates co-occur within host tissues (7, 8).

Among animal-associated microbes, bacteria belonging to the phylum Spirochaetes (spirochetes) are morphologically distinguished by their elongated, helically coiled or flat-wave cell shapes and their unique motility powered by periplasmic endoflagella residing between the inner and outer membranes (9, 10). This motility enables pathogenic spirochetes to navigate complex extracellular matrices, cross tissue barriers, and evade host immune responses (10, 11). Consequently, in medical and veterinary contexts, spirochetes are widely recognized as invasive pathogens causing multisystemic diseases, such as Lyme disease (*Borrelia burgdorferi*), syphilis (*Treponema pallidum*), leptospirosis (*Leptospira interrogans*), and severe periodontitis (*Treponema denticola*) (11–14). While these pathogens can transiently enter or translocate through host cells during tissue dissemination (15), eukaryotic intracellular environments present substantial barriers—including lysosomal degradation, cell-autonomous immune defenses, and oxidative stress—that typically lead to bacterial destruction or host cell death (7, 16). Within single-celled protists, such as wood-feeding termite gut flagellates, specific spirochetes have evolved obligate intracellular endosymbioses that provide key nutritional functions, including CO_2_-reductive acetogenesis and nitrogen fixation (17, 18). In contrast, across multicellular animals, spirochetes have hitherto been documented almost exclusively as extracellular epibionts attached to mucosal or epithelial surfaces (18, 19), or as transient invasive pathogens (11–15). Whether they can establish a stable, functional intracellular symbiosis within animal host cells has thus remained an open question.

Deep-sea hydrothermal vent and cold-seep mussels belonging to the family Mytilidae (subfamily Bathymodiolinae) serve as key models for chemosynthetic endosymbiosis, harboring dense populations of Gammaproteobacteria that utilize reduced chemicals, such as sulfide, methane, and hydrogen, to fuel host nutrition within specialized gill epithelial cells termed bacteriocytes (20, 21). Although the deep-sea vent mussel *Gigantidas horikoshii* has been recognized as a member of this group (22, 23), the molecular identity, metabolic capacity, and cellular localization of its symbionts have remained uncharacterized. Here, in resolving this uncharacterized symbiosis through a multi-scale approach combining metagenomics, metatranscriptomics, fluorescence *in situ* hybridization, transmission electron microscopy, and three-dimensional ultrastructural reconstruction via array tomography, we discovered that *G. horikoshii* harbors an intracellular micro-consortium within its bacteriocytes, comprising a primary sulfur-oxidizing Gammaproteobacterium (GhoSUP) and a co-resident spirochete (GhoSPI). Spatial and ultrastructural analyses show that GhoSPI co-localizes with GhoSUP along gill filaments, residing within the same closed, membrane-bound chambers inside bacteriocytes. Integrated genomic and transcriptomic analyses reveal active transcription of pathways for organic acid utilization, antioxidant defense, and host interactions in GhoSPI, providing functional support for this spatial co-existence. Based on these findings, we discuss the structural, cellular, genomic, and metabolic basis of cooperation between the chemosynthetic bacterium and the spirochete within this intracellular consortium.

## Results

### Genomic features and expression profiles reveal distinct metabolic roles of GhoSUP and GhoSPI in the gill symbiosis

Metagenomic sequencing was performed on *G. horikoshii* specimens, comprising the gill of one Sumisu Caldera individual and the whole-body samples of three Higashi-Aogashima juveniles (Table S1). Using these datasets, five metagenome-assembled genomes (MAGs) were reconstructed for two representative individuals, GhoSM1 and GhoHA4 (Table 1). Phylogenomic analysis placed GhoHA4SUP within a clade of sulfur-oxidizing Gammaproteobacteria predominantly comprising symbiotic bacteria, GhoHA4SPI within a clade of Spirochaetota mostly consisting of animal-associated bacteria, and GhoHA4CAM within Campylobacterota (Figure S1, Tables S2 and S3). Based on the high 16S rRNA gene identities (99.9% between GhoSM1SUP and GhoHA4SUP; 100% between GhoSM1SPI and GhoHA4SPI), these five MAGs were designated as three distinct symbiont lineages: GhoSUP, GhoSPI, and GhoCAM (Table 1). GhoSUP and GhoSPI occurred in both sample types, whereas GhoCAM was restricted to the whole-body samples of Higashi-Aogashima juveniles (Table 1). Metagenomic read mapping confirmed the consistent presence of MAGs GhoHA4SUP and GhoHA4SPI in the two additional Higashi-Aogashima juvenile specimens (GhoHA3 and GhoHA6), yielding high genome coverage (99.6% and 96.5% for GhoHA4SUP; 99.9% and 99.6% for GhoHA4SPI, respectively). In contrast, the lower abundance of GhoHA4CAM in these individuals resulted in lower or more variable genome coverage (27.2% and 68.5%, respectively) (Table S4). Genome-wide homology searches of predicted coding sequences showed that an average of 33.2% of the genes in the GhoSPI MAGs matched Spirochaetota, in contrast to the GhoSUP MAGs, where an average of 92.9% of genes matched Gammaproteobacteria or sulfur-oxidizing bacteria. Dual RNA-seq datasets were generated from the gills of two *G. horikoshii* individuals collected from the Higashi-Aogashima vent field, yielding 81.4–82.1 million paired-end reads per sample (Table S5), which were mapped to the three GhoHA4 MAGs to quantify gene expression levels as Z-scores. Although the GhoCAM genome encodes sulfur oxidation and carbon fixation via the reductive TCA cycle, reads mapping to GhoCAM were negligible in the gill transcriptomes (30–82 reads; 0.0% of clean reads) (Table S5), indicating its absence in the gills. Consequently, GhoCAM was excluded from subsequent analyses.

**Table 1.** Statistics on symbiont genomes of deep sea mussel *G. horikoshli*.

| MAGs | GhoSM1SUP | GhoSM1SPI | GhoHA4SUP | GhoHA4SPI | GhoHA4CAM |
|---|---|---|---|---|---|
| Host | GhoSM1 | GhoSM1 | GhoHA4 | GhoHA4 | GhoHA4 |
| Description | GhoSUP | GhoSPI | GhoSUP | GhoSPI | GhoCAM |
| Organism | Candidatus *Thiodubiliella* sp. | Spirochaetota bacterium | Candidatus *Thiodubiliella* sp. | Spirochaetota bacterium | *Sulfurovum* sp. |
| Number of contigs | 88 | 24 | 94 | 25 | 19 |
| Size (bp) | 1879681 | 2476425 | 1925721 | 2451628 | 2254233 |
| GC content (%) | 37.1 | 28.9 | 36.9 | 29 | 38.1 |
| Coverage | 1397 | 35 | 767 | 193 | 52 |
| Number of genes |  |  |  |  |  |
| CDS | 1652 | 2033 | 1697 | 1989 | 2337 |
| rRNA | 3 | 2 | 3 | 3 | 5 |
| tRNA | 41 | 35 | 38 | 35 | 37 |
| ncRNA | 3 | 2 | 3 | 2 | 2 |
| tmRNA | 1 | 1 | 1 | 1 | 1 |
| Accession | BAAMAR010000001-BAAMAR010000088 | BAAMAS010000001-BAAMAS010000024 | BAAMAT010000001-BAAMAT010000094 | BAAMAU010000001-BAAMAU010000025 | BAAMAV010000001-BAAMAV010000019 |

The GhoSUP genome encodes a complete cytoplasmic sulfur-oxidation pathway consisting of the dissimilatory sulfite reductase complex (*dsrABCEFH*), sulfate adenylyltransferase (*sat*), and adenosine-5′-phosphosulfate reductase (*aprABM*). Additionally, the genome contains an incomplete periplasmic Sox system (*soxABXYZ*) lacking soxCD. This truncated Sox system is predicted to drive the accumulation of elemental sulfur globules, which are subsequently oxidized in the cytoplasm via the Dsr complex (24). An encoded membrane-bound sulfide:quinone oxidoreductase (*sqr*) provides an entry point for sulfide into this sulfur-oxidation network (Table S6). Transcriptomic analysis confirmed active transcription of the cytoplasmic sulfur-oxidation pathway (*dsrABCEFH*, *aprABM*, and *sat*; Z-scores = 0.39 to 1.75), with the periplasmic Sox system exhibiting comparatively higher expression levels (Z-scores = 1.09 to 2.61) (Table S6). Beyond sulfur metabolism, GhoSUP encodes a membrane-bound nitrate reductase (*narGH*), showing lower expression levels (Z-scores = −0.16 to 0.31), whereas genes mediating hydrogen oxidation were not detected (Table S6). For carbon fixation, the genome encodes the Calvin-Benson-Bassham (CBB) cycle (including *rbcL*, *rbcS*, *and prkA*), and transcriptomic profiling confirmed its active expression (Table S6). Conversely, the TCA cycle was incomplete, lacking genes encoding citrate synthase (*gltA*), 2-oxoglutarate dehydrogenase complex (*sucAB*), succinate dehydrogenase (*sdhABC*), and malate dehydrogenase (*mdh*).

Unlike termite gut spirochetes (17), GhoSPI lacks genes required for nitrogen fixation and reductive acetogenesis (Table S7). Instead, it encodes diverse catabolic pathways for the utilization of short-chain organic acids, including propanoate, lactate, and acetate (Table S7). Propanoate is converted to succinate and pyruvate via the 2-methylcitrate cycle (*prpBCDE–acnA*, Z-score = 0.13 to 1.09) in a process linked to ATP generation. Lactate catabolism is similarly predicted to yield ATP via D-lactate dehydrogenase (*dld*) and the L-lactate dehydrogenase complex (*lldEFG*, Z-score = −1.42 to 1.06). Pyruvate derived from these pathways can be routed to acetate via acetyl-CoA, with ATP generated by either the Pta–Ack pathway (*pta*, *ackA*, and *acyP*) or acetyl-CoA synthetase (*acs*) (Z-score = 0.44 to 1.71). Additionally, the GhoSPI genome harbors complete pathways for glycolysis (Z-score = −0.70 to 1.45) and the tricarboxylic acid (TCA) cycle (Z-score = −0.96 to 2.19, including *sdhABC*, *sucABCD*, *icd*, and *mdh*; with *sucCD* at Z-score = 1.06 to 1.12), as well as F-type ATP synthase (*atpABCDEFGH*, with *atpE* exhibiting high expression at Z-score = 2.93) (Table S7). Because the genome lacks ubiquinol-cytochrome c reductase (*petABC*) for aerobic respiration, GhoSPI operates anaerobically, coupling these central carbon pathways to fermentation to regenerate NAD⁺.

To support its heterotrophic lifestyle, GhoSPI encodes a suite of nutrient importers (Table S8). These include transporters for sugars (*msmEFG*, Z-score = −1.60 to 0.70; *gtsABC*, Z-score = −1.57 to 2.34), peptides/nickel (ABC.PE.S, *oppA–F*, *yejA–F*, Z-score = −1.04 to 1.90), TRAP-type dicarboxylates (*dctPMQ*, Z-score = −1.61 to 2.06), and phosphate (*pstABCS*, Z-score = −0.12 to 2.29). Furthermore, genes mediating amino acid and polyamine catabolism (*speB*, *gabD*, *glyA*, and *serA*) were predominantly actively transcribed (Z-score = −1.01 to 1.53) (Table S7). For motility, GhoSPI encodes a complete flagellar system (basal body, hook, and motor), including two sheath proteins (*flaA*, Z-score = 0.42 to 0.48) and two core flagellins (*flaB*, Z-score = 1.01 to 2.06) (Table S7).

For potential host interactions, the GhoSPI genome encodes 119 DUF285 domain-containing proteins (Z-score = −1.10 to 2.26), eight β-propeller fold lactonases, predominantly with tandem domains (Z-score = 1.61 to 2.59), and four Fic-family effectors (with three of four actively transcribed at Z-score = 0.82 to 1.42) (Table S9). Proteins in the DUF285 domain-containing proteins and lactonase families were largely predicted to localize to the extracellular space or outer membrane (Table S9). Phylogenetic analyses of key host association proteins (ABC.PE.S, Fic effectors, and AhpC) showed that these proteins did not form monophyletic clades but clustered with diverse environmental, marine, or symbiotic bacteria (Figure S2). Furthermore, GhoSPI actively transcribes genes encoding a comprehensive ROS-scavenging system, including superoxide dismutase (*sod*, Z-score = 1.27), catalase (*katE*, Z-score = 1.75), alkyl hydroperoxide reductase (*ahpC*, Z-score = 0.15 to 1.20), and thioredoxin (*trxA*, Z-score = −0.09) (Table S7).

Additionally, to evaluate the symbiotic interface and the host’s physiological state in light of the metabolic capacities of the symbionts, we analyzed the transcriptomic profile of the host gill. The host gill highly expressed key genes involved in anaerobic energy metabolism, including phosphoenolpyruvate carboxykinase (*PCK*, Z-score = 3.44), malate dehydrogenases (*MDH1/2*, Z-score = 2.11 to 2.29), succinate dehydrogenase subunits (*SDHA–D*, Z-score = 1.76 to 4.13), propanoyl-CoA:succinate CoA transferase (*ASCT*, Z-score = 1.57), and propionyl-CoA carboxylase subunits (*PCCA/B*, Z-score = 1.43 to 1.50) (Table S10 and Figure S4).

### GhoSUP and GhoSPI co-localize intracellularly with distinct spatiotemporal colonization patterns

We analyzed the intracellular localization of GhoSUP and GhoSPI using FISH with specific 16S rRNA probes. Although glutaraldehyde fixation is unconventional for FISH, our technical improvements yielded higher-resolution images for Sumisu Caldera samples (Figure 1A) than conventional paraformaldehyde-fixed Higashi-Aogashima samples (Figures 1B–1E, Table S1). In the gill epithelium from both sites, GhoSUP was detected as numerous ∼1-µm spots clustered in the apical half of columnar cells, while the host nuclei occupied the basal side. GhoSPI co-localized in the same cells, showing a fibrous appearance that filled the spaces between GhoSUP spots, with a slightly higher density toward the extreme apical end (Figures 1A and 1B). In whole-body sagittal sections of Higashi-Aogashima juveniles, neither symbiont was detected in the posteriormost gill filaments. GhoSUP appeared in all subsequent anterior filaments, whereas GhoSPI was first detected slightly anterior to the onset of GhoSUP. GhoSPI was initially sparse in these transitional filaments but established stable co-localization with GhoSUP further anteriorly (Figure 1C). Outside the gills, both symbionts co-localized apically in a portion of the mantle (Figures 1D and 1E) but were absent in other organs.

**Figure 1.**
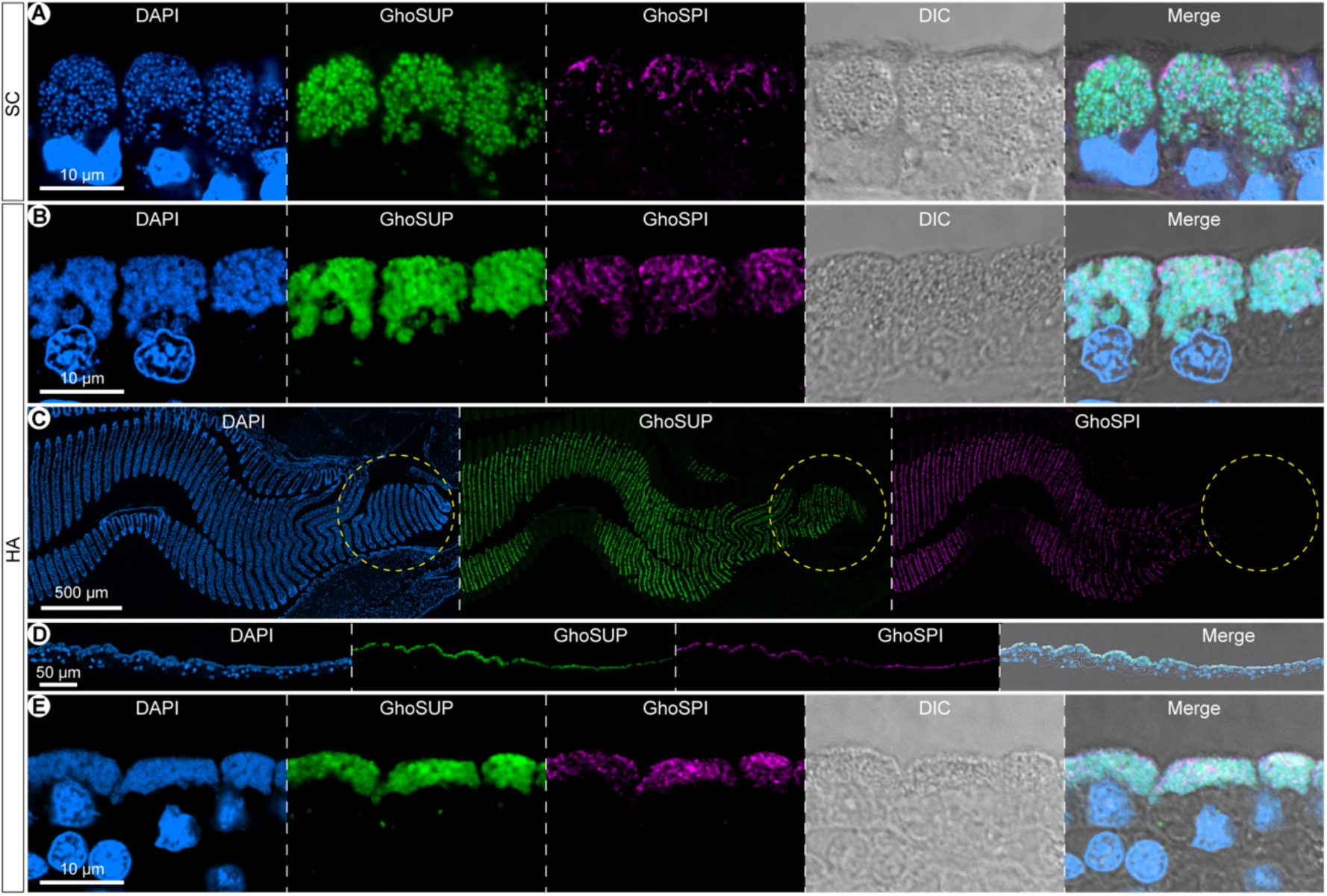
Spatiotemporal co-localization patterns of GhoSUP and GhoSPI. All panels show hybridization chain reaction-fluorescence in situ hybridization (HCR-FISH) images. Host nuclei and bacterial DNA are stained with DAPI (blue), while GhoSUP and GhoSPI are visualized with specific probes in green and magenta, respectively. DIC indicates differential interference contrast image. (A) High-resolution image of gill epithelial cells from a Sumisu Caldera (SC) specimen fixed with glutaraldehyde. The apical side is at the top. (B–E) Images of a juvenile specimen collected from the Higashi-Aogashima vent field (HA) and fixed with paraformaldehyde. (B) High-magnification image of the gill filaments. The apical side is at the top. (C) Low-magnification image of the gill. The dorsal side is at the top, and the posterior side is on the right. Note that within the yellow dotted circle indicating the posterior end of the gill, DAPI and GhoSUP signals are present, but GhoSPI is not detected. (D) Low-magnification image of the mantle. (E) High-magnification image of the mantle, demonstrating the concurrent intracellular presence of both symbionts.

### The two symbionts co-reside within closed, interconnected host chambers

Transmission electron microscopy (TEM, Table S1) of the inner gill epithelium revealed alternating columnar cells and elongated intermediate cells (25), the latter characterized by dense apical microvilli and basal nuclei (Figure 2A). The columnar cells housed apical, circular-to-oval bacteria-like structures (∼0.5–1.0 µm in diameter) identified as GhoSUP based on morphology (20) (Figures 2B and 2C). These bacteria exhibited nucleoids, distinct double membranes, and electron-dense periplasmic particles (∼0.1–0.2 µm in diameter) (Figure 2C). All these GhoSUP cells were localized within the membranous chamber structures of the host cell, with multiple bacteria present within each chamber compartment, and some compartments appeared to be interconnected (Figures 2A and 2B). Within the same chambers, elongated bacteria-like structures (∼0.2– 0.3 µm thick) were identified as GhoSPI based on the FISH results and the characteristic ultrastructure of spirochetes (10) (Figures 2B, 2D, and 2E). GhoSPI cells were often in close contact with the chamber wall and spanned multiple chamber compartments (Figures 2D and 2E). In addition, electron-dense concentric multilamellar structures were observed between the symbiotic chambers and the basal nucleus (Figures 2A and 2F). Similarly, in the samples from Higashi-Aogashima, in addition to gill epithelial cells, mantle epithelial cells housed both GhoSUP and GhoSPI within shared intracellular chambers (Figures 2G and 2H).

**Figure 2.**
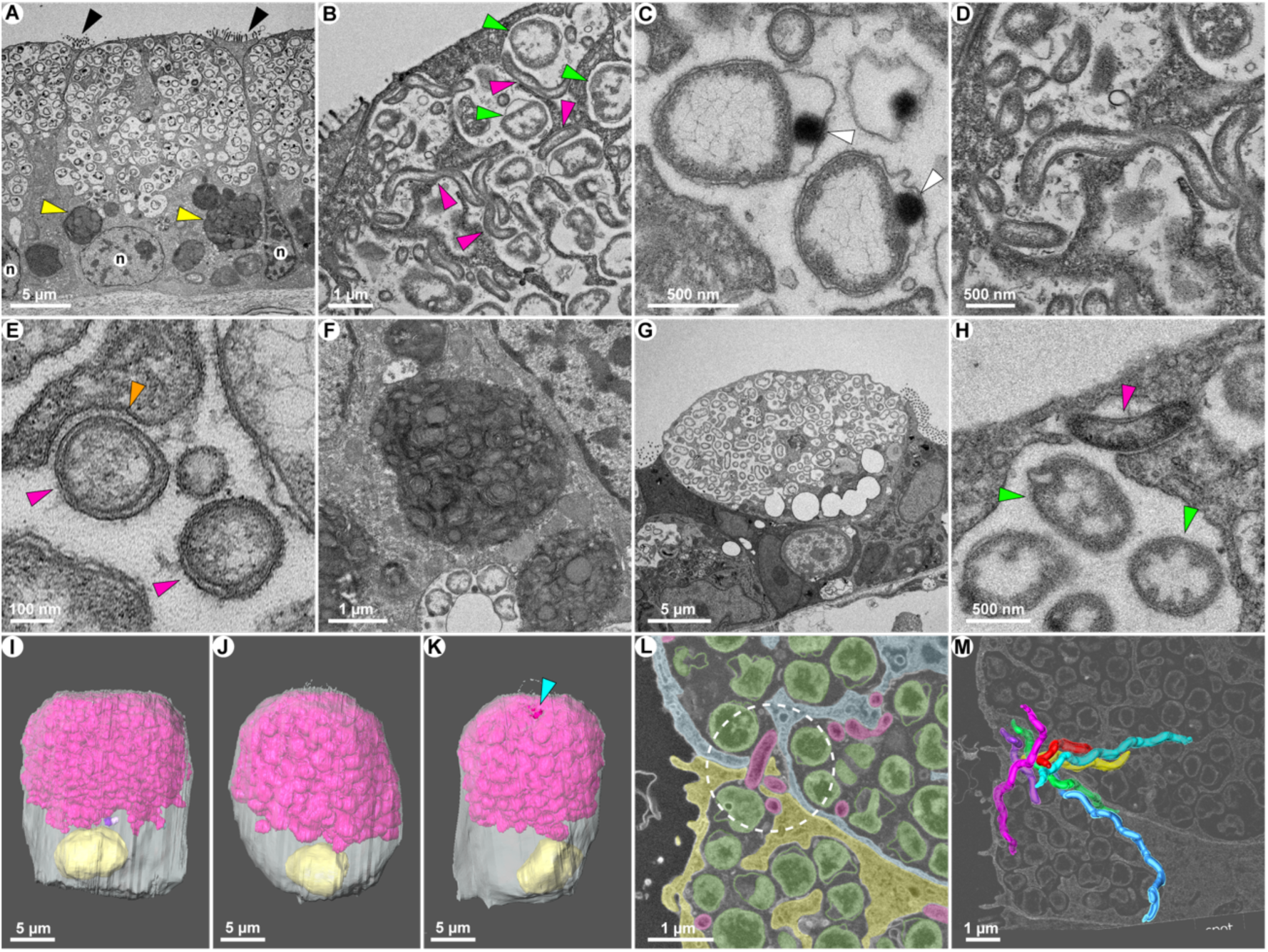
Ultrastructural architecture and 3D reconstruction of bacteriocytes and co-resident symbionts. (A–H) Representative transmission electron microscopy (TEM) images. (A–F) Gill epithelial cells from a Sumisu Caldera specimen. (A) Overview of a bacteriocyte housing bacteria within chambers, flanked by intermediate cells with dense apical microvilli (black arrowheads). The apical side is at the top. Bacteria-enclosing chambers are located apically, while regions of highly electron-dense concentric multilamellar structures are visible on the basal side (yellow arrowheads). n: nucleus. (B) Close-up of the apical side of the bacteriocyte. Sulfur-oxidizing bacteria (GhoSUP, green arrowheads) and spirochetes (GhoSPI, magenta arrowheads), identified by their morphology, co-reside within the same chamber. (C) Close-up of sulfur-oxidizing bacteria showing clear inner and outer membranes with electron-dense granular structures in the periplasm (white arrowheads). (D) Close-up of a spirochete extending across adjacent chamber compartments. (E) Cross-section of a spirochete (magenta arrowheads) demonstrating distinct inner and outer membranes, with the host chamber membrane closely apposed to a part of its outer surface (orange arrowhead). (F) Close-up of the highly electron-dense concentric multilamellar structure on the basal side of the bacteriocyte. (G, H) Bacteria-harboring mantle cells from a Higashi-Aogashima specimen. (G) Overview of a mantle cell. (H) Close-up of the mantle cell showing that, similar to the gills, sulfur-oxidizing bacteria (green arrowheads) and spirochetes (magenta arrowheads) co-reside within the same chamber. (I–M) 3D reconstruction of gill bacteriocytes using array tomography. (I– K) 3D reconstructions of three entire bacteriocytes (I: cell 1, J: cell 2, and K: cell 3). The interconnected chambers are shown in different colors, with the largest chamber in magenta. The nucleus and cytoplasm are labeled with yellow and slightly transparent white, respectively. The apical side is at the top. In cell 3 (K), the light blue arrowhead indicates the site of penetration into an adjacent cell created by the extending spirochetes. See Supplemental Data for 3D rotation movies. (L) Close-up of cell 3 showing spirochetes extending across two adjacent cells. The cytoplasm of the two host cells is colored blue and yellow, respectively. Sulfur-oxidizing bacteria and spirochetes are shown in green and magenta, respectively. Note the spirochete within the white dotted circle extending across the boundary between the two cells. See Supplemental Data for an animation of the serial sections. The apical side is on the left. (M) 3D reconstruction of seven spirochete cells extending across two host cells in cell 3, overlaid with a representative cross-sectional image indicating the positions of the two bacteriocytes. The apical side is on the left. See Supplemental Data for a 3D animation.

Furthermore, three-dimensional reconstruction via array tomography (cells 1–3) revealed that the symbiotic chambers were highly interconnected (99.6–100% volume, Table S11) and lacked apical openings to the external environment (Figures 2I–2K and Supplemental Movies 1–3). However, in one examined cell, a lateral chamber opening allowed GhoSPI cells to extend into a neighboring host cell’s chamber (Figures 2K–2M and Supplemental Movies 4 and 5).

## Discussion

This study demonstrates that an intracellular spirochete (GhoSPI) integrates into a chemosynthetic micro-consortium with a sulfur-oxidizing Gammaproteobacterium (GhoSUP) within interconnected and enclosed intracellular chambers in the gill bacteriocytes of the deep-sea hydrothermal vent mussel *Gigantidas horikoshii*. Multi-omics data indicate that GhoSUP autotrophically drives the symbiotic system via active sulfur oxidation (Figure S3). Our 3D reconstruction revealed that the chambers encompassing the symbionts are isolated from the extracellular environment (Figures 2I–2K; Supplemental Movies 1–3), in contrast to the related vent mussel *Bathymodiolus septemdierum*, which possesses open, interconnected symbiotic chambers (26). This closed system, alongside basal concentric multilamellar structures (Figures 2A and 2F), indicative of intracellular digestion (25, 27), suggests that the host sequesters GhoSUP, potentially facilitating the harvesting of synthesized nutrients.

Within these isolated chambers, GhoSPI exhibits a heterotrophic lifestyle (Figure S3). Lacking reductive acetogenesis and nitrogen fixation genes typical of termite-gut spirochetes (17, 18), GhoSPI transcribes various transporters, amino acid catabolic enzymes, and TCA/ATP synthesis genes (e.g., *atpE*; Z-score = 2.93) (Tables S7 and S8), indicating reliance on host- or GhoSUP-derived organics. Our metatranscriptomic data suggest that GhoSPI may function as an intracellular environmental stabilizer. The host actively transcribes anaerobic energy metabolism genes (e.g., *PCK*, *MDH*, *SDH*, *ASCT*, and *PCCA/B*) (Table S10 and Figure S4), likely reflecting hypoxia induced by the enclosed chamber and GhoSUP’s oxygen consumption (28). Consistent with anaerobic pathways characterized in other bivalves (29), this host metabolic shift likely leads to the accumulation of short-chain organic acids (e.g., succinate, acetate, and propionate) within the chambers. While the autotrophic GhoSUP lacks the 2-methylcitrate cycle for propionate assimilation (Table S6), GhoSPI actively transcribes genes for the transport and utilization of these acids (*dctPMQ*, Z-score = −1.61 to 2.06; *acs*, Z-score = 1.13; and *prpBCDE*, Z-score = 0.13 to 1.09) (Tables S7 and S8). This contrast suggests that GhoSPI utilizes these host-derived metabolic byproducts as carbon and energy sources, which could mitigate local acidosis and contribute to stabilizing the host’s cellular microenvironment.

Oxidative stress management represents another critical challenge within this enclosed intracellular environment. Physiological stress associated with deep-sea environmental fluctuations is known to elevate reactive oxygen species (ROS) levels and trigger antioxidant enzyme responses in related *Bathymodiolus* gills (30). Our metatranscriptomic profiling revealed distinct antioxidant strategies between the two coexisting symbionts. Heterotrophic GhoSPI actively transcribed key antioxidant genes, including superoxide dismutase (*sod*, Z-score = 1.27), catalase (*katE*, Z-score = 1.75), and peroxiredoxin (*ahpC*, Z-score = 0.15 to 1.20). In contrast, while the autotrophic GhoSUP showed low transcription of *katE* (Z-score = −1.49) and *ahpC* (Z-score = −0.44), it actively transcribed a glutathione-dependent peroxiredoxin (*pgdX*, Z-score = 1.54) (Tables S6 and S7). This suggests that GhoSUP and GhoSPI deploy distinct yet complementary antioxidant enzyme systems to cope with oxidative stress within the host cellular microenvironment.

In contrast to GhoSUP, where an average of 92.9% of genes matched Gammaproteobacteria or sulfur-oxidizing bacteria, only 33.2% of GhoSPI genes showed top hits to Spirochaetota. While this pattern may partly reflect the unique phylogenetic position of GhoSPI, it also suggests a potentially chimeric genomic architecture. In particular, phylogenetic analyses of representative key factors for host interaction and cellular homeostasis—Fic effectors (31, 32), peptide transporters (ABC.PE.S) (33), and peroxiredoxin (AhpC) (34, 35)—revealed polyphyletic distributions rather than monophyletic spirochete clades (Figure S2). This suggests that these molecular tools were acquired via multiple horizontal gene transfers from diverse environmental, marine, or symbiotic bacteria. Consequently, parallel acquisitions likely played a major role in expanding the repertoire of these gene families in GhoSPI. Together, these acquired toolkits, including AhpC and ABC.PE.S, likely equip GhoSPI with the capabilities required for cellular homeostasis and stress mitigation during intracellular coexistence. In addition, physical adhesion and tissue colonization may be supported by the highly expanded DUF285 domain-containing proteins, which show homology with BspA-family proteins (36) from other host-associated bacteria (37, 38). Certain DUF285-containing proteins are known to directly modulate host immune responses and cellular signaling (36). Transcriptomic data demonstrated that GhoSPI actively transcribes these DUF285 proteins, alongside Fic-family effectors and lactonases (Table S9). Collectively, these functions—including DUF285-mediated interactions, host signaling manipulation by Fic effectors (31, 32), and metabolite modification by lactonases (39, 40)—likely constitute a multifaceted host interaction machinery.

Beyond these toolkits, GhoSPI exhibits dynamic spatial behavior. FISH observations showed a preferential apical localization of GhoSPI in bacteriocytes (Figures 1A and 1B), and 3D reconstruction demonstrated that GhoSPI cells can extend across host cell boundaries into the chambers of adjacent cells (Figures 2K–2M). Supporting this active motility, transcriptomic data revealed high expression of flagellar genes, particularly those encoding the core flagellin (*flaB*) (Table S7). Confined within enclosed chambers without direct seawater access, flagellar propulsion, supported by high flagellin expression, may provide the physical force that enables GhoSPI to navigate restrictive host cellular spaces and maintain its apical positioning, thereby escaping basal lysosomal degradation. Such active motility could also facilitate host tissue colonization and subsequent transcellular migration (15, 41).

Spatiotemporal colonization dynamics further support GhoSPI’s role as an environmental stabilizer. Since deep-sea mussel gills grow continuously from the posterior end (42), the anteroposterior axis reflects chronological development. Accordingly, FISH observations revealed that initial GhoSUP colonization in newly formed posterior filaments was followed by GhoSPI integration in more mature anterior regions, leading to consistent co-localization (Figure 1C). This lag suggests a chronological sequence in which GhoSUP acts as a pioneer, establishing initial colonization and chemolithoautotrophy. Vigorous sulfur oxidation and oxygen consumption likely elevate oxidative stress (30) and induce hypoxia (28) within enclosed chambers. Following this environmental shift, GhoSPI integrates at later stages, utilizing active motility, ROS-scavenging, and organic acid assimilation to help mitigate localized oxidative stress and acidosis. This complementary functional relationship likely optimizes resource allocation, allowing GhoSUP to dedicate energy primarily to high-output chemolithoautotrophy. Furthermore, the concurrent intracellular presence of GhoSUP and GhoSPI in juvenile mantle epithelial cells (Figures 1D, 1E, 2G, and 2H) probably reflects a transient extra-branchial infection phase typical of early-stage deep-sea mussels (43, 44), underscoring the tight spatial linkage between these symbionts. Thus, these observations suggest that GhoSPI plays an important role in establishing and maintaining a stable intracellular micro-consortium, rather than a coincidental co-infection. However, while active motility provides physical force, the molecular mechanisms underlying the initial invasion of GhoSPI into bacteriocytes and enclosed chambers remain to be investigated.

In conclusion, our findings demonstrate that an intracellular spirochete (GhoSPI) can integrate as a stable, functional partner within animal cells in a deep-sea chemosynthetic symbiosis. Within this enclosed microenvironment, this spirochete appears to function as an intracellular environmental stabilizer, utilizing metabolic and host-interaction toolkits putatively derived from diverse origins to mitigate localized acidosis and oxidative stress, thereby facilitating stable intracellular coexistence. Beyond this specific deep-sea symbiosis, these findings provide a new perspective on the functional landscape of animal-microbe interactions, shedding light on mechanisms linking mutualistic endosymbiosis and intracellular infections.

## Materials and Methods

### Animal sampling

*Gigantidas horikoshii* was collected from hydrothermal vent sites in the Sumisu Caldera, Izu-Ogasawara Arc, Japan, at a depth of 694 m during cruise YK24-13C (19–27 August 2023) using the human-occupied vehicle (HOV) *Shinkai 6500* operated by the R/V *Yokosuka* (Japan Agency of Marine-Earth Science and Technology; JAMSTEC), and on the Higashi-Aogashima vent field, Izu-Ogasawara Arc, Japan, at depths of 746 and 759 m during cruise KM24-03 Leg 1 (16–25 April 2024) using the R/V *Kaimei*-operated *KM-ROV* (JAMSTEC). Mussels collected from the Sumisu Caldera were immediately bisected sagittally upon retrieval by cutting the adductor muscles with a scalpel, and then fixed in 2.5% glutaraldehyde (GA) in seawater filtered with a 0.2-μm filter unit (Nalgene, Rochester, NY, USA) (filtered seawater, FSW) at 4°C. Some of the mussels collected from the Higashi-Aogashima vent field were immediately bisected sagittally with a razor upon retrieval, with a small portion of mantle tissue excised and frozen in liquid nitrogen, and the remaining part was fixed in either 2.5% GA in FSW or 4% paraformaldehyde (PFA) in 1 × phosphate-buffered saline (PBS). Samples fixed with GA were stored as-is at 4°C, while those fixed with PFA overnight were stepwise dehydrated with an ethanol series and stored at −30°C. The remaining specimens collected from Aogashima were frozen in liquid nitrogen and stored at −80°C.

### Species identification

DNA was extracted from the partially excised and frozen mantle specimens collected from Higashi-Aogashima using the DNeasy Blood and Tissue Kit (Qiagen, Hilden, Germany) or NucleoSpin Tissue XS (Machery-Nagel, Düren, Germany) according to the manufacturer’s instructions. In addition, the gills excised from mussels that had been collected from the Sumisu Caldera and fixed in GA were beaten with zirconia beads using a TissueLyzer II (Qiagen) (25 Hz, 2 s, 2 times). DNA was extracted from the bead-treated suspensions according to the manufacturers’ protocol. PCR was conducted using primer sets designed for the bivalve mitochondrial COI gene (LCO1490 (45) and H691 (46)) and PrimeSTAR HS polymerase (Takara, Ohtsu, Japan). The amplicons were sequenced by the Sanger method with the same primers employed for amplification, and mussel species identification was performed via a BLASTN search against the NCBI database (http://blast.ncbi.nlm.nih.gov/Blast.cgi).

### Metagenome sequencing and assembly

DNA was extracted from the whole bodies of three frozen juvenile individuals collected from Higashi-Aogashima using the DNeasy Blood & Tissue Kit (Qiagen, Hilden, Germany). Additionally, DNA was extracted from the GA-fixed gills of a single individual collected from the Sumisu Caldera. Metagenome libraries were constructed using the KAPA HyperPrep Kit for Illumina platforms (Roche, MA, USA) according to the manufacturer’s protocol. Paired-end sequencing was performed on the Element Biosciences AVITI platform (CA, USA) with a 2 × 300 bp read length. Raw sequence reads were sequentially processed using Trimmomatic v0.39 (47) to trim adapter and low-quality sequences. Cleaned reads were assembled using CLC Genomics Workbench v26 (Qiagen) with the following parameters: 64 bp word size, 500 bp bubble size, and mapping reads back to the contigs with a 0.9 length fraction and a 0.9 similarity fraction. The metagenome contigs were binned and reassembled using an R-script-based pipeline (https://github.com/MadsAlbertsen/multi-metagenome) as previously described (48). Briefly, binning was performed based on differences in GC content, tetranucleotide frequency, and sequencing coverage across the contigs. Extension of the targeted contigs was performed using the PriceTI assembler (49). Finally, the generated contigs were inspected and manually corrected. Completeness and contamination of the draft genomes were calculated using CheckM (50) using the lineage-specific marker sets for Gammaproteobacteria, Spirochaetota, and Campylobacterota. Raw metagenomic sequencing reads were deposited in the DDBJ Sequence Read Archive (DRA; https://www.ddbj.nig.ac.jp/dra/index-e.html) under accession numbers DRR1085880 (GhoSM1) and DRR1085882 (GhoHA4), and the assembled genome sequences were deposited at DDBJ/EMBL/GenBank under accession numbers BAAMAR010000000 (GhoSM1SUP), BAAMAS010000000 (GhoSM1SPI), BAAMAT010000000 (GhoHA4SUP), BAAMAU010000000 (GhoHA4SPI), and BAAMAV010000000 (GhoHA4CAM), under BioProject accession PRJDB42931.

### Bioinformatic analysis

The coding sequences in the symbiont genomes were identified using Prodigal ver. 2.6.3(51). Their functional annotations were assigned using the Kyoto Encyclopedia of Genes and Genomes (KEGG) orthology database (52) as a reference. Noncoding RNA genes, such as rRNAs and tRNAs, were predicted using Infernal ver. 1.1.5 (53) and tRNAscan-SE ver. 2.0.12 (54). Average nucleotide identity (ANI) and average amino acid identity (AAI) were estimated using OrthoANI ver. 1.20 (55) and CompareM ver. 0.1.2 (https://github.com/dparks1134/CompareM), respectively. Subcellular localizations of proteins were predicted using PSORTb v3.0 (56), CELLO v2.5 (57), and DeepLocPro 1.0 (58).

### Metatranscriptome analysis

Total RNA was extracted from PFA-fixed gills of juveniles collected from the Higashi-Aogashima vent field using the RNeasy FFPE Kit (Qiagen) as follows. A portion of the gill (approximately six filaments) was excised from two juveniles and transferred to a sample tube. Buffer PKD (240 µL) was added along with zirconia beads, and the tissue was disrupted using a TissueLyzer II (Qiagen) at 25 Hz for 2 s, twice. Proteinase K (10 µL) was added, and the mixture was incubated at 56 °C for 3 h. The subsequent steps were performed according to the manufacturer’s protocol, and RNA was eluted in 14 µL of RNase-free water.

Total RNA was treated with the RiboCop rRNA Depletion Kit for Bacteria (Lexogen Inc., Vienna, Austria) to obtain ribosomal RNA (rRNA)-depleted RNA. The quantity and quality of the RNA were assessed using the Qubit RNA HS Assay Kit with a Qubit 1.0 Fluorometer (Thermo Fisher Scientific, Waltham, MA, USA) and the Agilent DNF-472 (15 nt) HS RNA Kit with an Agilent 5200 Fragment Analyzer System (Agilent Technologies, Santa Clara, CA, USA). Strand-specific RNA-seq libraries were constructed via cDNA synthesis using the SMART-Seq Stranded Kit (Clontech, Mountain View, CA, USA). The resulting libraries were sequenced on the AVITI platform (Element Biosciences, CA, USA) with a 2 × 150 bp read length.

To estimate the expression levels of symbiont genes, RNA-seq reads were processed using the Trimmomatic program and then mapped to the genome of each MAG using Bowtie2 v2.5.4 (59). Read counts assigned to annotated genes were quantified using featureCounts (60). Transcripts per million (TPM) values were calculated using the Python script RNAnorm (https://github.com/genialis/RNAnorm).

To estimate host gene expression levels, de novo transcriptome assembly was performed using Trinity v2.15.2 (61) with default parameters. All assembled transcript sequences were further processed to obtain a non-redundant transcript set using the EvidentialGene tr2aacds pipeline (http://arthropods.eugenes.org/EvidentialGene/). This pipeline selects the “best” set of de novo assembled transcripts based on coding potential. Protein-coding regions were predicted using the TransDecoder program (https://github.com/TransDecoder/TransDecoder). Functional annotation was performed via homology searches against the National Center for Biotechnology Information (NCBI) non-redundant (nr) protein database and the Kyoto Encyclopedia of Genes and Genomes (KEGG) database (62) with an E-value threshold of 1 × 10^−5^. The predicted protein-coding transcripts were aligned against the nr database using DIAMOND v2.1.12 (63) in the BLASTx mode with an E-value threshold of 1 × 10^−5^. BUSCO v6.1.0 (64) was used to evaluate the completeness of the *G. horikoshii* transcriptome assembly. Table S5 summarizes the metatranscriptomic sequencing statistics and the results of the BUSCO completeness evaluation. Host gene expression levels were quantified using the RNA-Seq by Expectation-Maximization (RSEM) software v1.2.23 (65) with Bowtie2.

To determine representative expression levels for both symbiont and host genes, TPM values for each gene were log2-transformed after adding a pseudocount of 1 [log2(TPM + 1)]. The arithmetic mean of these log-transformed values was then calculated across the biological replicates. Where indicated, mean values were back-transformed to the original TPM scale (2^mean − 1) to represent the geometric mean. Subsequently, Z-scores were calculated using the formula Z = (Xg − Mg) / Sd, where Xg is the mean log2-transformed expression value for a given gene, and Mg and Sd are the overall mean and standard deviation across all genes, respectively. This Z-score represents the relative rank of a given gene’s expression within the transcriptome-wide distribution. Raw metatranscriptomic sequencing reads were deposited in the DDBJ Sequence Read Archive (DRA) under accession numbers DRR1085884 (GhoHA12) and DRR1085885 (GhoHA15) under BioProject accession PRJDB42931.

### Phylogenomic analysis and taxonomic classification

Taxonomic classification of the MAGs was performed using the classify_wf workflow in GTDB-Tk v2.5.2 (66) based on the Genome Taxonomy Database (GTDB) release r220 (67). To reconstruct phylogenomic trees, the amino acid sequences of single-copy marker genes (Table S2) were extracted from the predicted proteomes of the target genomes and their related reference genomes (Table S3). Multiple sequence alignment was performed using MAFFT v7.312 (68), and ambiguously aligned positions were removed using trimAl v1.2 (69). After concatenating the alignments, maximum likelihood (ML) phylogenomic trees were inferred using IQ-TREE 2 v2.3.5 (70). Best-fit amino acid substitution models were determined using ModelFinder integrated within IQ-TREE based on the Bayesian Information Criterion (BIC): Q.plant+F+I+R3 for sulfur-oxidizing bacteria, LG+F+R4 for Spirochaetota, and LG+F+R3 for Campylobacterota. Branch support was assessed using the ultrafast bootstrap approximation with 2,000 replicates.

### Phylogenetic analysis for identification of representative key factors

To investigate the evolutionary origins of representative key factors implicated in GhoSPI’s intracellular adaptation (i.e., Fic effectors, the peptide/nickel transporter ABC.PE.S, and the peroxiredoxin AhpC), local BLASTP searches were performed against the translated CDSs of the symbiont genomes with an *E*-value cutoff of 1e−5, using well-characterized sequences as queries. For Fic effectors, the candidate sequences were further analyzed using the InterPro server (71) to confirm the presence of the Fido domain (IPR003812) and sequence alignments to verify the conservation of the functional HPFxxGNGR core motif (31). Subsequently, multiple sequence alignments were generated using only the extracted core Fido domains (97–125 amino acids). As for ABC.PE.S and AhpC, the full-length amino acid sequences encoded by the identified genes were used without domain trimming. To infer their phylogenetic relationships, reference sequences from diverse bacterial taxa were retrieved from the NCBI (https://www.ncbi.nlm.nih.gov/) non-redundant protein database. Death-on-curing (Doc) family toxins, OppA, and thioredoxin-dependent thiol peroxidase sequences were selected as outgroups for Fic effectors, ABC.PE.S, and AhpC, respectively. Multiple sequence alignments with the ClustalW algorithm and subsequent ML phylogenetic tree reconstructions were performed using MEGA12 (72). s were reconstructed using MEGA12. The best-fit amino acid substitution models determined and applied for the analyses were WAG+G+I for Fic effectors and AhpC, and LG+G+F for ABC.PE.S. Branch support was assessed using 1,000 bootstrap replicates.

### Fluorescent in situ hybridization (FISH)

In this study, due to constraints on the available samples, several special procedures were required to perform the FISH analysis. Samples collected from the Sumisu Caldera and fixed with GA, as well as specimens collected from the Higashi-Aogashima vent field, fixed with PFA, and preserved in ethanol, were used. To enhance the fluorescence intensity, the hybridization chain reaction (HCR)-FISH method (73, 74) was applied with the following modifications. A piece of gill was excised from individuals fixed with GA and washed with PBS. For PFA-fixed juveniles, the shells were removed from one-half of the bisected mussels. These specimens were dehydrated stepwise using an ethanol series, embedded in paraffin, and sectioned to a thickness of 4 µm. The sections were dewaxed in xylene and stepwise rehydrated with ethanol and PBS. The sections were treated with 1 mg/ml sodium borohydride in PBS on ice for 5 min twice. After washing with PBS, the sections were treated with 3% hydrogen peroxide in PBS at room temperature and washed with PBS. The sections were then treated with 1 µg/ml proteinase K in PBS at 37°C for 20 min, followed by post-fixation with 4% paraformaldehyde in PBS. The pre-hybridization solution, which contained 20 mM Tris-HCl, 0.9 M NaCl, 0.01% SDS, 20% formamide, 10% sodium dextran sulfate, and 1% blocking reagent (Roche, Basel, Switzerland), was applied to the samples, covered with Parafilm (Amcor, Zurich, Switzerland), and incubated in a moist chamber at 46 °C for 15 min. Hybridization was then carried out using a hybridization solution containing 0.5 µM of two initiator probes in a moist chamber at 46 °C for 2 h. One probe (5 ′ - TACGCCCTAAGAATCCGAACCCTATGAAAAAGGTGAGCGCCCAGATAAGCTA-3′) was specifically designed for the 16S rRNA of sulfur-oxidizing symbiotic bacteria in several deep-sea mussel species, including *G. horikoshii* (allowing for two-base mismatches), with an iR sequence (75) attached to the 5′ end. The other probe (5′ - CCAGTTATCAGTAGTCCGTCCTTCATTTTTTGCAGATAGCTAATCATTACTCAC C-3′) was specifically designed for the 16S rRNA of the spirochete identified in *G. horikoshii* in this study (remaining exclusively specific to this spirochete even when allowing for up to two base mismatches), with an iC sequence (75) attached to the 5′ end. After hybridization, the excess probe was washed away in a wash solution containing 225 mM NaCl, 20 mM Tris-HCl pH 7.5, and 0.01% SDS at 48°C for 15 min, twice. The sections were incubated in 1 M NH_4_Cl solution at room temperature for 15 min, followed by quenching autofluorescence with TiYO (Nepagene, Ichikawa, Japan) for 1 h. During the quenching, the R1/C1 and R2/C2 amplifier probes (5 µM each; R1 and R2 probes were labeled with Alexa Fluor 647, and C1 and C2 probes were labeled with Alexa Fluor 555) were incubated in amplification buffer, which contained 0.05 M Na_2_HPO_4_, 0.9 M NaCl, 10% sodium dextran sulfate, 1 × Denhardt’s Solution (Nakarai Tesque, Kyoto, Japan), 0.1 mg/ml heparin (Sigma-Aldrich, St. Louis, MO, USA), 0.1 mg/ml torula yeast RNA (Sigma-Aldrich), 0.1% Tween 20, 0.01% SDS, at 95 °C for 1.5 min and then 30 min at 25°C.

The sections were immersed in amplification buffer at room temperature for 15 min, and a mixture of R1/C1 and R2/C2 amplifier probes was applied to them. After incubation at 46°C for 2 h, the sections were washed with phosphate-buffered saline (PBS) containing 0.1% Tween 20 for 10 min at 4°C and counterstained with DAPI (2 µg/ml) in PBS. Finally, the sections were mounted with ProLong Glass (Thermo Fisher Scientific) and left at 4°C overnight or longer to solidify. FISH images were captured using a Leica Stellaris STED system (Leica Microsystems, Wetzlar, Germany) and processed using Adobe Photoshop 2026 (Adobe Systems, San Jose, CA, USA).

### Electron microscopy and array tomography

The preparation and observation of specimens for transmission electron microscopy (TEM) were performed using the gills of mussels that had been collected from the Sumisu Caldera and Higashi-Aogashima vent field and fixed with GA, as previously described (76).

Scanning electron microscopy (SEM) observations for three-dimensional reconstruction (array tomography) of bacteriocytes were conducted as previously described (26, 77). A gill sample from one individual collected from the Sumisu Caldera, which had been embedded in Epon 812 (TAAB, Aldermaston, UK) for TEM, as described above, was used. A total of 300 serial ultrathin sections, each 80 nm thick, were prepared and collected onto a coverslip. The sections were stained with 2% uranyl acetate, coated with osmium, and observed under an FEI Quanta 450FEG scanning electron microscope (Thermo Fisher Scientific, Waltham, MA, USA) operated at 5 kV. For three bacteriocytes, images were captured at 15,000 × magnification, covering the entire cell thickness, yielding 219, 239, and 249 serial section images, respectively. Additionally, to observe the localization of spirochetes spanning two adjacent bacteriocytes, 26 consecutive images were acquired at 45,000 × magnification.

Images of the serial sections were aligned into a single stack using AMIRA v6.3 (Thermo Fisher Scientific). Structures of interest (symbiotic chambers, cell nuclei, cell membranes, and bacterial cells) were manually traced for segmentation throughout the image stack using the Microscopy Image Browser (MIB) software v24.1 (78). After segmentation, the data were returned to AMIRA, and 3D tomographic images were constructed using post-processing, including surface rendering and smoothing. In addition, the volumes of the segmented structures were calculated using the surface area-volume function of AMIRA.

## Acknowledgments

We are grateful to Yasuo Furushima (JAMSTEC), Chong Chen (JAMSTEC), the captains and crew of the R/V *Kaimei* and R/V *Yokosuka*, the operation teams of ROV *KM-ROV* and HOV *Shinkai 6500*, and other researchers who were aboard the research cruises for their assistance with sample collection. Cruise YK24-13C (chief scientist: Shinsuke Kawagucci, JAMSTEC) was supported by the Council for Science, Technology and Innovation (CSTI), Cross-ministerial Strategic Innovation Promotion Program (SIP), and National Platform for Innovative Ocean Developments (lead agency: JAMSTEC).

## Author contributions

Conceptualization, T.I. and A.Y.; Methodology, T.I.; Investigation, T.I., Y.T., A.Y., Y.Y., and M.T.; Data curation, T.I. and Y.T.; Visualization, T.I.; Writing – original draft, T.I.; Writing – review & editing, T.I., Y.T., and A.Y.

