## Supplemental Figure S1_4 for "Intracellular spirochete integrates into a chemosynthetic micro-consortium of a deep-sea vent mussel"

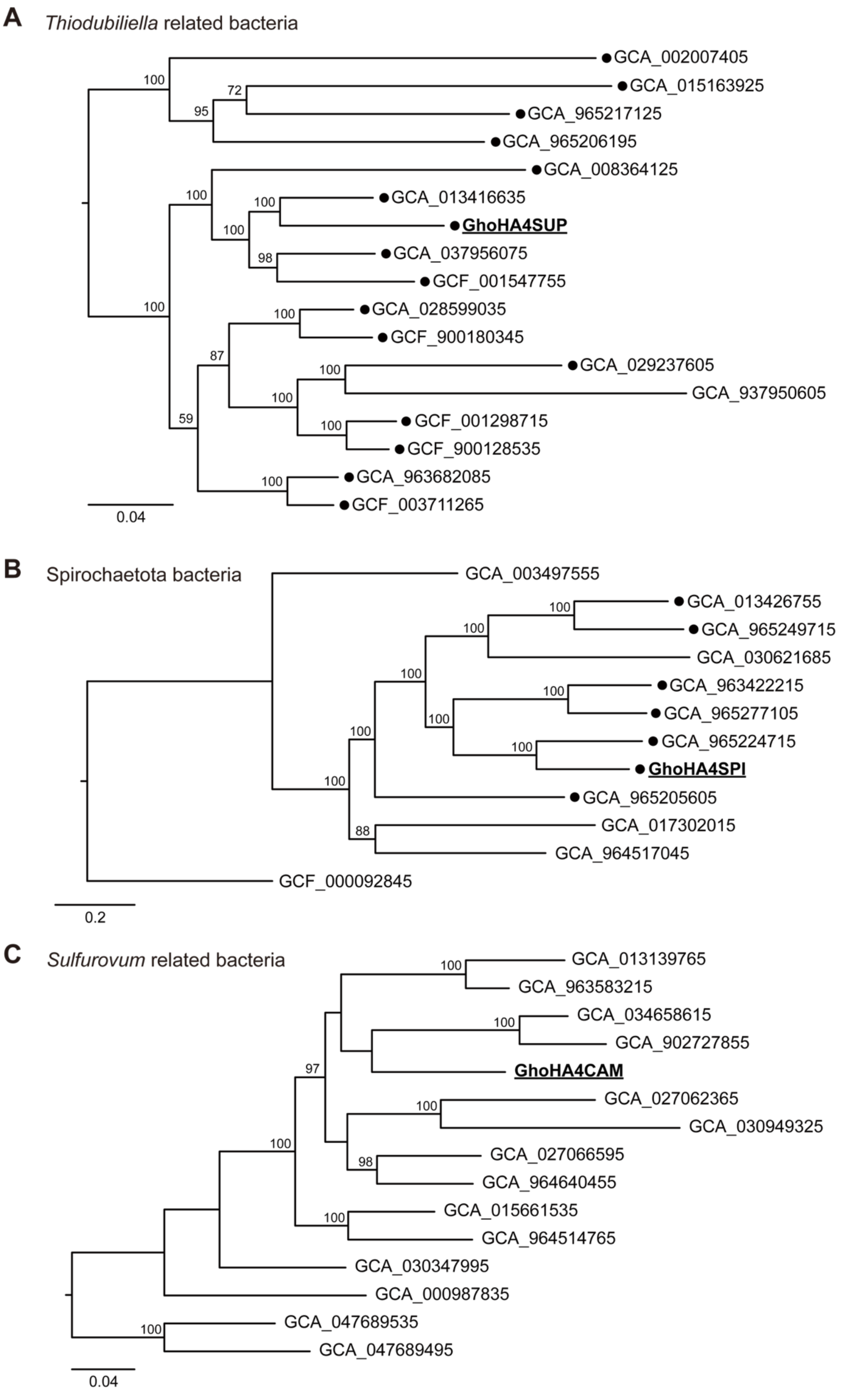
A
Thiodubiliella related bacteria
GCA_002007405
GCA_015163925
GCA_965217125
GCA_965206195
GCA_008364125
GCA_013416635
GhoHA4SUP
GCA_037956075
GCF_001547755
GCA_028599035
GCF_900180345
GCA_029237605
GCA_937950605
GCF_001298715
GCF_900128535
GCA_963682085
GCF_003711265
0.04
B
Spirochaetota bacteria
GCA_003497555
GCA_013426755
GCA_965249715
GCA_030621685
GCA_963422215
GCA_965277105
GCA_965224715
GhoHA4SPI
GCA_965205605
GCA_017302015
GCA_964517045
GCF_000092845
0.2
C
Sulfurovum related bacteria
GCA_013139765
GCA_963583215
GCA_034658615
GCA_902727855
GhoHA4CAM
GCA_027062365
GCA_030949325
GCA_027066595
GCA_964640455
GCA_015661535
GCA_964514765
GCA_030347995
GCA_000987835
GCA_047689535
GCA_047689495
0.04

**Figure S1. Phylogenetic placement of three bacterial lineages recovered from *Gigantidas horikoshii*.**

Maximum-likelihood phylogenomic trees were constructed based on conserved genes for (A) Thiodubiliella-related sulfur-oxidizing bacteria, (B) Spirochaetota, and (C) Sulfurovum-related Campylobacterota. Black dots indicate strains reported as symbionts in (A) or animal associates in (B). The conserved genes and taxa used in each tree are listed in Tables S2 and S3, respectively. Node support was evaluated using ultrafast bootstrap analysis with 2,000 replicates. The scale bars represent amino acid substitutions per site.

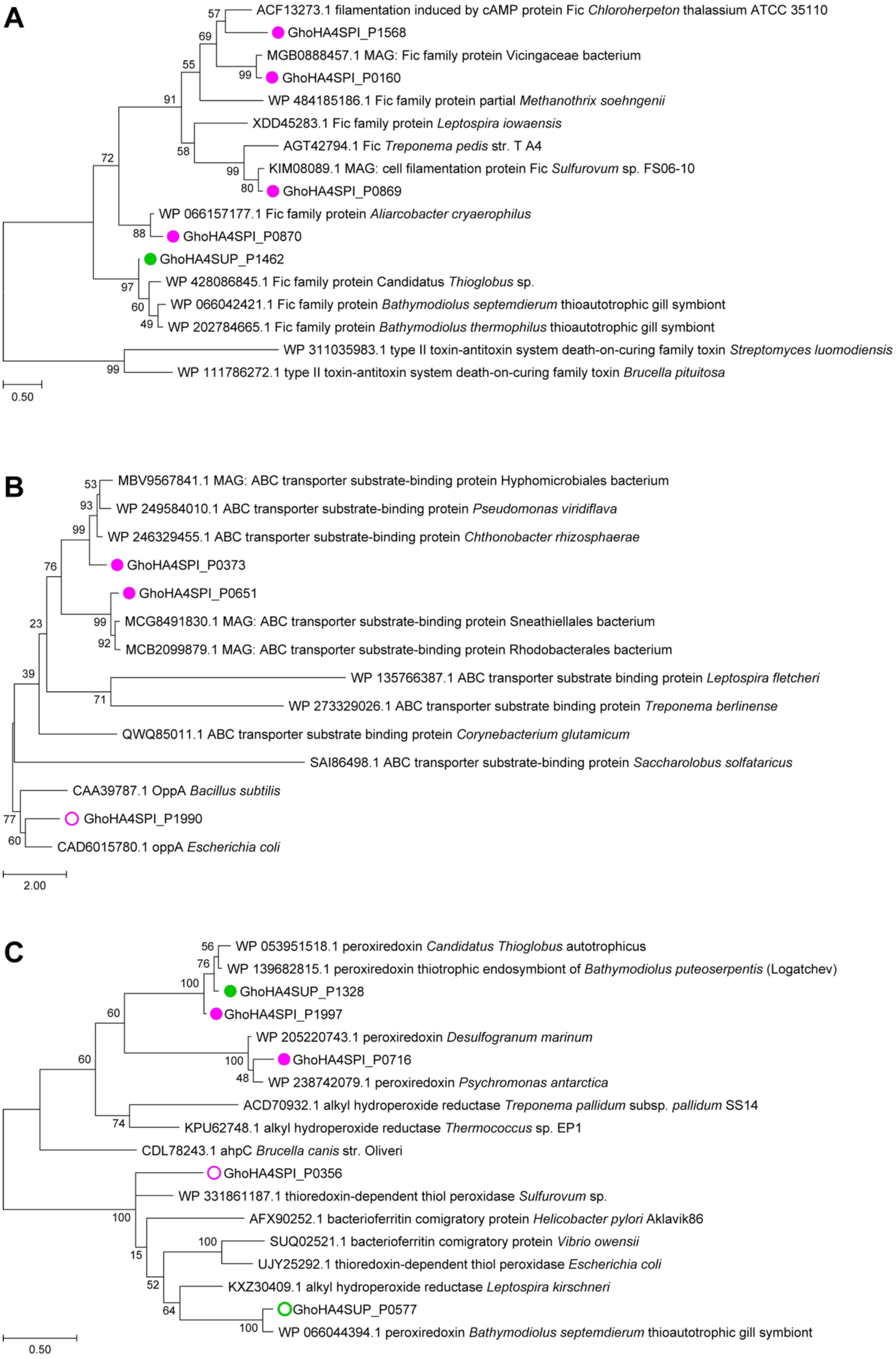
A
ACF13273.1 filamentation induced by cAMP protein Fic Chloroherpeton thalassium ATCC 35110
GhoHA4SPI_P1568
MGB0888457.1 MAG: Fic family protein Vicingaceae bacterium
GhoHA4SPI_P0160
WP 484185186.1 Fic family protein partial Methanothrix soehngenii
XDD45283.1 Fic family protein Leptospira iowaensis
AGT42794.1 Fic Treponema pedis str. T A4
KIM08089.1 MAG: cell filamentation protein Fic Sulfurovum sp. FS06-10
GhoHA4SPI_P0869
WP 066157177.1 Fic family protein Aliarcobacter cryaerophilus
GhoHA4SPI_P0870
GhoHA4SUP_P1462
WP 428086845.1 Fic family protein Candidatus Thioglobus sp.
WP 066042421.1 Fic family protein Bathymodiolus septemdierum thioautotrophic gill symbiont
WP 202784665.1 Fic family protein Bathymodiolus thermophilus thioautotrophic gill symbiont
WP 311035983.1 type II toxin-antitoxin system death-on-curing family toxin Streptomyces luomodiensis
WP 111786272.1 type II toxin-antitoxin system death-on-curing family toxin Brucella pituitosa
57
69
55
99
91
72
58
99
80
88
97
60
49
99
0.50
B
MBV9567841.1 MAG: ABC transporter substrate-binding protein Hyphomicrobiales bacterium
WP 249584010.1 ABC transporter substrate-binding protein Pseudomonas viridiflava
WP 246329455.1 ABC transporter substrate-binding protein Chthonobacter rhizosphaerae
GhoHA4SPI_P0373
GhoHA4SPI_P0651
MCG8491830.1 MAG: ABC transporter substrate-binding protein Sneathiellales bacterium
MCB2099879.1 MAG: ABC transporter substrate-binding protein Rhodobacterales bacterium
WP 135766387.1 ABC transporter substrate binding protein Leptospira fletcheri
WP 273329026.1 ABC transporter substrate binding protein Treponema berlinense
QWQ85011.1 ABC transporter substrate binding protein Corynebacterium glutamicum
SAI86498.1 ABC transporter substrate-binding protein Saccharolobus solfataricus
CAA39787.1 OppA Bacillus subtilis
GhoHA4SPI_P1990
CAD6015780.1 oppA Escherichia coli
53
93
99
76
99
92
23
39
71
77
60
2.00
C
WP 053951518.1 peroxiredoxin Candidatus Thioglobus autotrophicus
WP 139682815.1 peroxiredoxin thiotrophic endosymbiont of Bathymodiolus puteoserpentis (Logatchev)
GhoHA4SUP_P1328
GhoHA4SPI_P1997
WP 205220743.1 peroxiredoxin Desulfogranum marinum
GhoHA4SPI_P0716
WP 238742079.1 peroxiredoxin Psychromonas antarctica
ACD70932.1 alkyl hydroperoxide reductase Treponema pallidum subsp. pallidum SS14
KPU62748.1 alkyl hydroperoxide reductase Thermococcus sp. EP1
CDL78243.1 ahpC Brucella canis str. Oliveri
GhoHA4SPI_P0356
WP 331861187.1 thioredoxin-dependent thiol peroxidase Sulfurovum sp.
AFX90252.1 bacterioferritin comigratory protein Helicobacter pylori Aklavik86
SUQ02521.1 bacterioferritin comigratory protein Vibrio owensii
UJY25292.1 thioredoxin-dependent thiol peroxidase Escherichia coli
KXZ30409.1 alkyl hydroperoxide reductase Leptospira kirschneri
GhoHA4SUP_P0577
WP 066044394.1 peroxiredoxin Bathymodiolus septemdierum thioautotrophic gill symbiont
56
76
100
60
60
100
48
74
100
15
100
52
64
100
0.50

**Figure S2. Maximum-likelihood phylogenetic trees of representative key factors implicated in the intracellular adaptation of GhoSPI.**

Phylogenetic relationships of **(A)** Fic effectors, **(B)** the peptide/nickel transporter ABC.PE.S, and **(C)** the peroxiredoxin AhpC. The bootstrap values (1,000 replicates) are shown at the nodes. The scale bar represents amino acid substitutions per site. For Fic effectors (A), the tree was generated using extracted core Fido domains (97–125 amino acids), and death-on-curing (Doc) family toxins were used as the outgroup. For ABC.PE.S (B) and AhpC (C), full-length amino acid sequences were used, and OppA and thioredoxin-dependent thiol peroxidase sequences were selected as outgroups for ABC.PE.S and AhpC, respectively. Gene IDs derived from GhoSPI and GhoSUP are marked with filled magenta and green dots, respectively. Gene IDs from GhoSPI and GhoSUP that clustered within the outgroup clades are indicated by open magenta and open green dots, respectively.

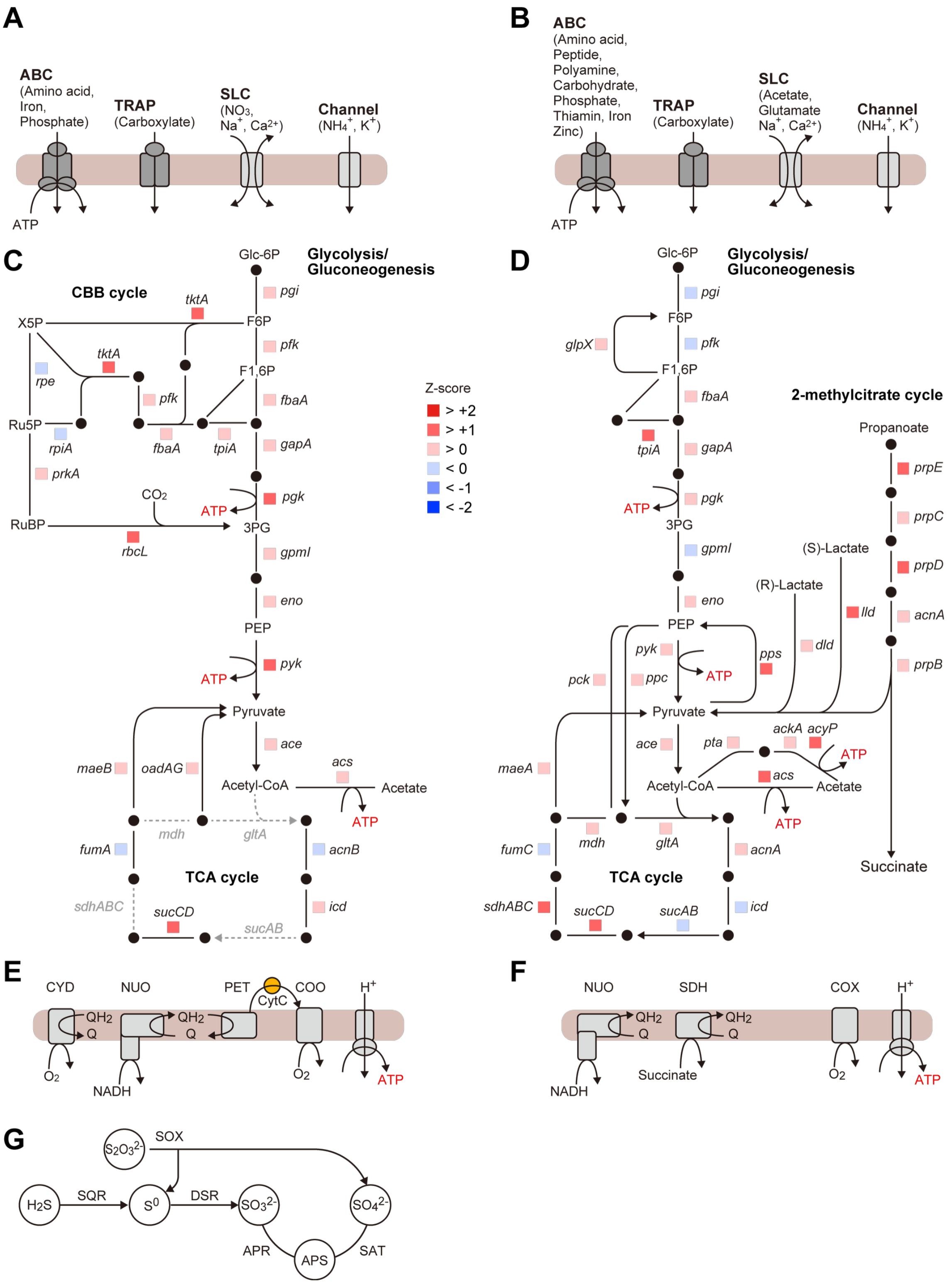


**Figure S3. Genome-derived map of predicted metabolic pathways and transport pathways in GhoSUP and GhoSPI.**

(A and B) Membrane transport systems and substrate uptake mechanisms in GhoSUP (A) and GhoSPI (B). Transporters and ion channels embedded in the cell membrane (brown band) are shown as solid blocks, with arrows indicating transport directions. (C and D) Central carbon metabolic pathways in GhoSUP (C) and GhoSPI (D). Arrows denote reaction directions, with adjacent colored boxes representing relative gene expression levels based on Z-scores. Expression levels were classified by Z-score (based on the standard normal distribution): $Z > 0$ indicates expression above the mean (top 50%); $Z > 0.5$ corresponds to the top 30.9% (i.e., moderately high expression); $Z > 1$ corresponds to the top 15.9% (i.e., high expression); and $Z > 2$ corresponds to the top 2.3% (i.e., very high expression). Dashed lines indicate defective genes. (E and F) Aerobic respiratory chains and membrane energy transduction in GhoSUP (E) and GhoSPI (F). (G) Sulfur oxidation pathway, present exclusively in GhoSUP. Gene abbreviations and expression values are provided in Tables S6 and S7, and transporter details in Table S8. Abbreviations: ABC, ATP-binding cassette transporter; TRAP, tripartite ATP-independent periplasmic transporter; SLC, solute carrier; CBB, Calvin–Benson–Bassham; NUO, NADH-quinone oxidoreductase complex; NQR, $Na^+$-transporting NADH:ubiquinone oxidoreductase; PET, ubiquinol-cytochrome c reductase; CCO, cbb3-type cytochrome c oxidase; CYD, cytochrome *bd* ubiquinol oxidase; COX, cytochrome c oxidase; SDH, succinate dehydrogenase; CytC, reduced cytochrome c; ATPase, F-type $H^+$-transporting ATPase; SOX, sulfur oxidation protein complex; SQR, sulfide:quinone oxidoreductase; DSR, reverse dissimilatory sulfite reductase; APR, adenylylsulfate reductase; SAT, sulfate adenylyltransferase.

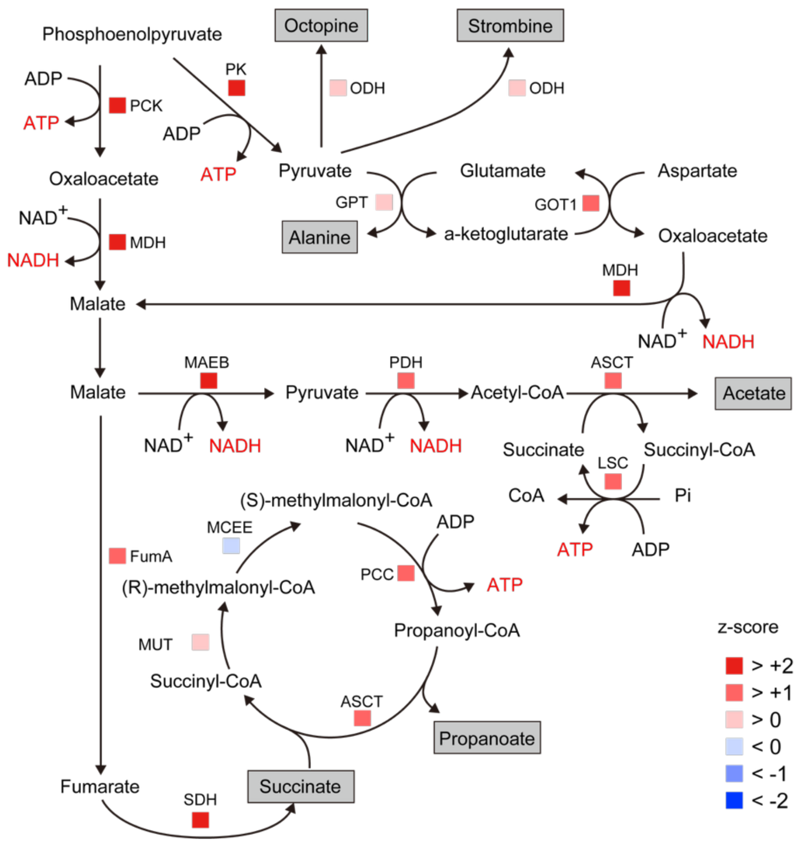


**Figure S4. Transcriptome-derived map of predicted anaerobic energy metabolism in the host animal.**

Under hypoxic or anaerobic conditions, the host mussel *Gigantidas horikoshii* utilizes anaerobic energy pathways, producing metabolites such as alanine, opines, succinate, acetate, and propanoate. During sustained anaerobiosis, acetate and propanoate are synthesized via mitochondrial pathways to support continued ATP generation. Abbreviations for the associated genes are provided in Table S10. Relative gene expression levels are indicated by box colors corresponding to Z-scores.
